# Gene regulatory networks and essential transcription factors for de novo originated genes

**DOI:** 10.1101/2024.12.19.629391

**Authors:** Junhui Peng, Bing-Jun Wang, Nicolas Svetec, Li Zhao

## Abstract

The regulation of gene expression is crucial for the functional integration of evolutionarily young genes, particularly those that emerge de novo. However, the regulatory programs governing the expression of de novo genes remain unknown. To address this, we applied computational methods to single-cell RNA sequencing data, identifying key transcription factors likely instrumental in regulating de novo genes. We found that transcription factors do not have the same propensity for regulating de novo genes; some transcription factors contain more de novo genes than others in their regulon. Leveraging genetic and genomic tools in *Drosophila*, we further examined the role of two key transcription factors and the regulatory architecture of novel genes. Our findings identify key transcription factors associated with the expression of de novo genes and provide new insights into how modifications in existing transcription factors enable the emergence, maintenance, and regulation of de novo genes.

## Introduction

Gene expression is orchestrated by a complex network of regulatory machinery, including but not limited to transcription factors (TFs), enhancers, repressors, and 3D chromosomal organization ^1,2^. This precise control of expression is essential to ensure specific gene activation at the right time and location, enabling the spatial and temporal control required for cellular differentiation, development, and adaptation to environmental stimuli. A key challenge in evolutionary biology is understanding how novel gene expression is integrated into these networks, particularly how regulatory systems adapt to incorporate accurate regulation of new genes and enable their proper functions or buffer their possible detrimental effects ^3^.

New genes emerge through several mechanisms, such as gene duplication, retrotransposition, horizontal gene transfer, and de novo gene birth ^4^. Once formed, these genes must be incorporated into existing regulatory networks to perform various biological functions. This integration, in return, requires a specific expression that may involve the emergence of new enhancers and the modification of TF activity. However, until now, it is unknown how changes in TF activities globally or specifically regulate the expression of new genes, especially new genes without preexisting regulatory elements, such as de novo genes and retroposed genes.

TFs, which bind to specific DNA sequences to regulate gene expression, are critical in defining the context in which genes are expressed. In particular, changes in TFs have been shown to expand regulatory networks ^5^. The changes in the copy number of TFs also may show a large effect on downstream expressional regulation through changes in TF expression or the binding affinity of specific DNA sequences ^5,6^. In simple terms, there are two possible mechanisms by which TFs may bind to de novo genes. First, de novo genes may acquire binding sites in their promoter or enhancer regions that can be recognized and regulated by a particular TF. Alternatively, TFs may alter their binding affinity in specific tissues to regulate a broader or different set of genes, including de novo genes. While both mechanisms are plausible, it remains unclear whether one or both contribute to the expression of de novo genes.

*Drosophila melanogaster* is an ideal model for investigating the dynamics of gene regulation because of the well-characterized genome and advanced genetic tools. The evolution of new genes, especially duplicated genes and potential de novo-originated genes, is overall well-characterized. Most of the evolutionarily young genes are enriched in the testis, presumably because of the role of sexual selection in the spread and fixation of new genes. In previous work, we used single-cell RNA sequencing (scRNA-Seq) to provide an unprecedented resolution of novel gene expression ^7,8^. We observed that very young de novo-originated genes, including those that are still polymorphic in populations ^9^, show specific spatio-temporal expression patterns in germ cells, suggesting that within a short time frame, the regulatory machinery is able to regulate these new genes precisely and accurately. However, it is unclear how this is achieved. How do TFs regulate new gene expression? Are there key TFs that specifically contribute to the regulation of novel genes? How does the emergence of novel binding sites *in cis* and potential changes in binding affinity of transcription factors *in trans* orchestrate accurate gene expression?

We use the *Drosophila* testis as a model to address this question. In addition to the enrichment of young genes in this tissue ^9,10^, the germ cell types in the testis are relatively simple, consisting of only a few major cell types involved in spermatogenesis ^7,11^. This simplicity facilitates the dissection of complex regulatory functions. Moreover, the number of transcription factors regulating young genes is comparatively small ^12^, enabling the identification of key transcription factors that may have a significant impact on the regulation of novel genes.

In this study, we use scRNAseq to identify regulons associated with novel gene expression in *D. melanogaster* testis. We identified potential key transcription factors linked to new gene expression. Using *Drosophila* genetics and -omics approaches, we illuminate the regulatory machinery underlying de novo gene expression, offering insights into how regulatory networks evolve to accommodate new gene functions and innovations in gene regulation.

## Results

### De novo genes have higher cell-type/tissue specificity and are expressed in a variety of tissues and cells

To gain a deeper understanding of the expression patterns of de novo gene candidates across major tissues and cell types, we used the Fly Cell Atlas dataset ^13^ to characterize the expression pattern for each of the 555 de novo gene candidates inferred from our previous work^14^. In the Fly Cell Atlas dataset, we were able to detect the expression of 498 out of the 555 identified de novo gene candidates and compute their scaled expression levels ^7^. The scaled expression describes how much the expression of a gene in a specific tissue or cell type deviates from the mean expression (Material and Methods). From the scaled expression of the 498 de novo gene candidates, we found that most of the de novo gene candidates are tissue- and cell-type-biased (Figure 1A and 1B), with 466 (86%) of them being tissue-biased and 475 (88%) being cell-type-biased, while the proportions for other annotated protein-coding genes are only 59% and 63%, respectively (supplementary data 1, Figure 1C and 1D). The top five enriched tissues correspond to 399 (80%) tissue biased de novo gene candidates, including 278 (55.8%) in the testis, 72 (14.5%) in the accessory glands, 20 (4%) in the fat body, 11 (2.2%) in the female reproductive tract, 9 (1.8%) in the head, and 9 (1.8%) in the gut. The top five enriched cell types correspond to 389 (78%) cell-type-biased de novo gene candidates, including 262 (52.6%) in male germline cells, 75 (15.1%) in accessory gland cells, 25 (5%) in fat cells, 16 (3.2%) in male- and 11 (2.2%) in female- reproductive system cells. There are also several de novo gene candidates that show biased expression in tissues such as antenna (8, 1.6%) and trachea (4, 0.8%), and cells such as sensory neurons (7, 1.4%) and hemocytes (7, 1.4%). In addition, we found that even at a finer cell type scale, where the 17 major cell types were further categorized into 250 distinct types, some de novo gene candidates showed clustered expression in some specific cell types (Figure S1). Overall, the biased tissues or cell types play key roles in mating, reproductive, sensory, and immune responses, suggesting possible important functions for de novo genes in these tissues.

**Figure 1.**
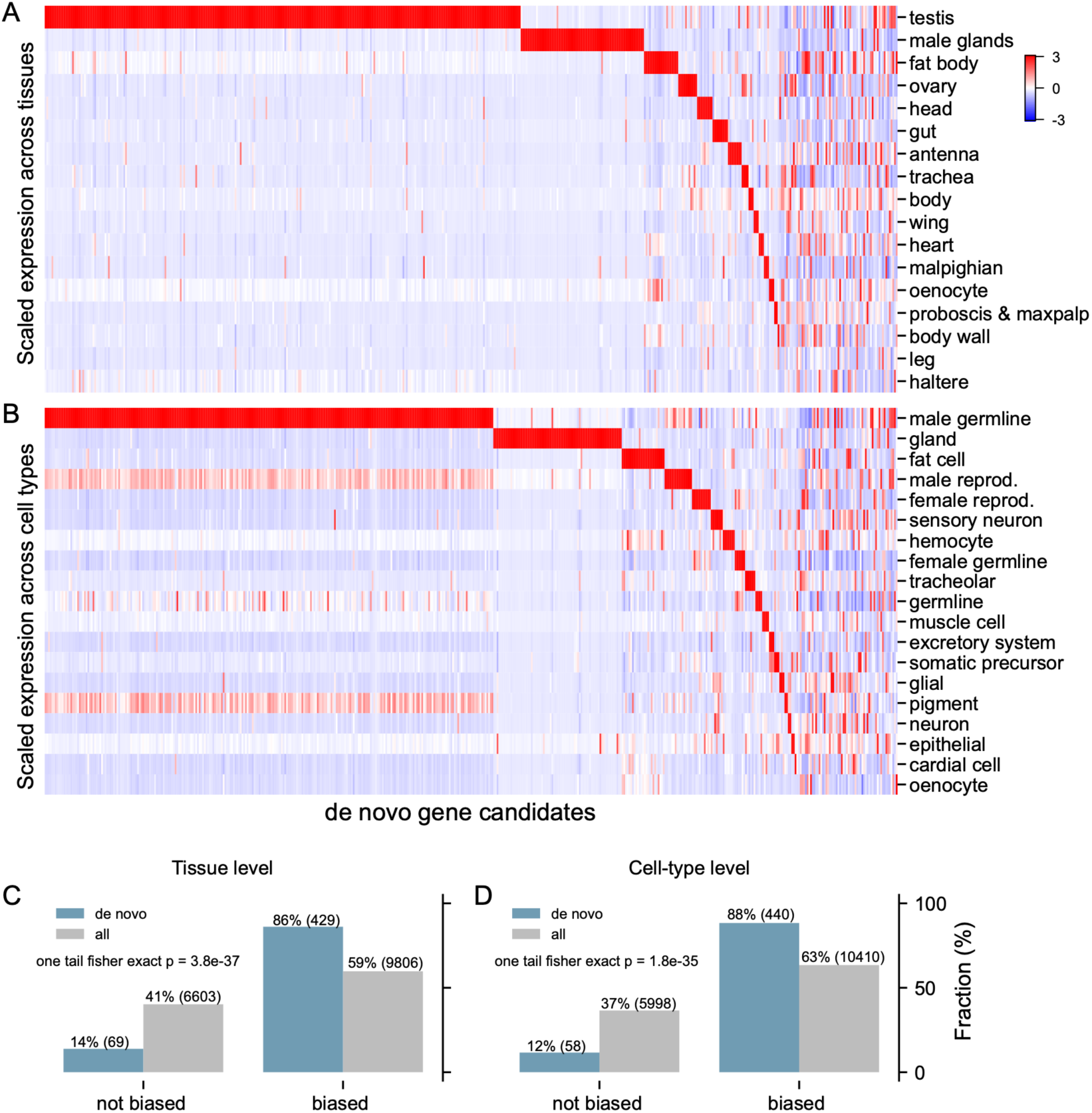
Expression patterns of de novo gene candidates across different tissues and major cell types. A) Scaled expression of de novo genes across different tissues. The tissues are ranked by the number of de novo genes biasedly expressed. B) Scaled expression of de novo genes across different major cell types. The major cell types are ranked by the number of de novo genes showing biased expression. The fractions of tissue- (C) and cell type- (D) biased de novo genes (colored in gray) and other annotated protein-coding genes (colored in blue) are also shown. The corresponding numbers of genes showing biased expression are labeled, indicating that de novo gene candidates are more likely to show biased expression in both tissue level (one-tailed Fisher’s exact test p = 3.8e-37) and cell-type level (one-tailed Fisher’s exact test p = 1.8e-35).

### Many de novo genes are regulated by only a few regulons

To understand the regulatory machinery underlying de novo genes in D. melanogaster, we carried out SCENIC analysis to find out if there are any master transcription factors that regulate the expression of de novo genes. To do this, we built an expression matrix including only de novo gene candidates, transcription factors, as well as other testis-biased genes inferred from FlyAtlas2 ^15^ as control. We further filtered out cells where no de novo genes were expressed and transcription factors were lowly expressed (Material and Methods). The SCENIC prediction process was similar to Li et al. ^13^. Interestingly, the predictions showed that the expression of most de novo gene candidates (60%, 300 of 498) were regulated by only 83 transcription factors (Figure 2A, supplementary data 2), which are less than 10 percent of the total transcription factors analyzed. Unlike the de novo gene candidates that showed highly biased expression patterns (Figure 1), the 83 transcription factors showed less tissue-specific patterns (Figure 2B), suggesting that de novo gene expression bias does not simply derive from an expression bias of their transcription factors. Although the TFs show different tissue specificity, their expression among different tissues have significantly higher correlations with their predicted targets compared to other non-target genes (Figure S2). Among the 83 transcription factors, 31 were predicted to regulate the expression of 10 or more de novo gene candidates, which is in line with the observation that de novo genes are enriched in a few tissues or cell types and is also consistent with our hypothesis that there might be key TFs regulating de novo genes. The top 31 transcription factors showed slightly but not significantly higher tissue specificity, with 15 showing no tissue specificity, four testis specific, three midgut specific, one hindgut specific, one accessory gland specific, one brain specific, one ovary specific, one thoracicoabdominal ganglion specific, and one salivary gland specific (Table S1). Interestingly, many of the de novo genes were predicted to be regulated by two or more TFs and some TFs were themselves predicted to be regulated by others, exhibiting a complex transcriptional regulatory network (Figure S3, S4).

**Figure 2.**
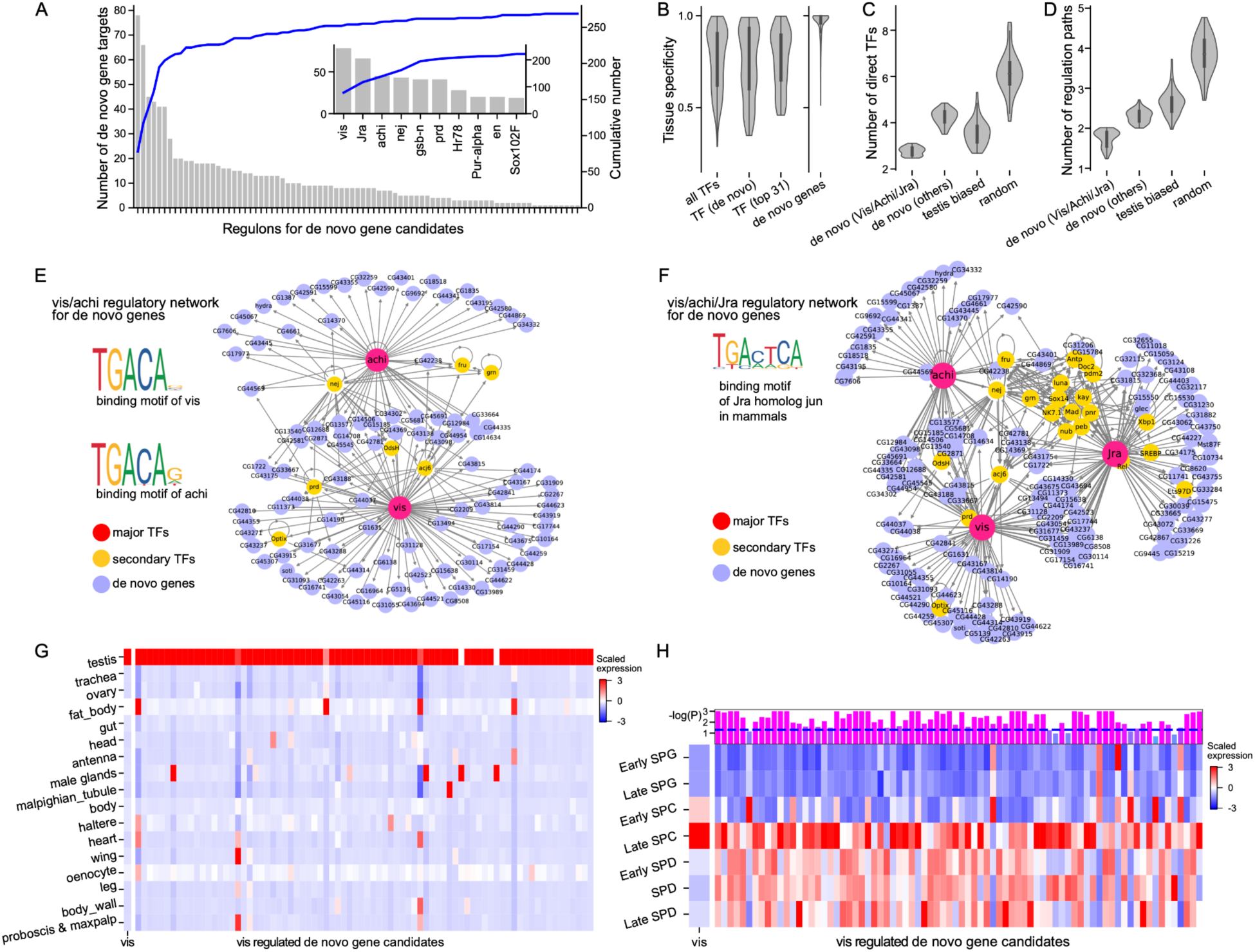
Transcriptional regulation of de novo genes. (A) Most de novo genes in D. melanogaster were predicted to be regulated by only a few transcription factors. The top three transcription factors are Vis, Jra, and Achi. (B) The transcription factors did not show high tissue specificity compared to de novo genes. De novo genes that are predicted to be regulated by the top three transcription factors (Vis/Achi/Jra), which we denoted as VAJ-regulated de novo genes or de novo (Vis/Achi/Jra) show more directed transcription regulations, with less number of direct transcription factors (C) and less number of transcriptional regulation paths. A detailed description of the two metrics can be found in Material and Methods. (E) The transcriptional regulatory network of Vis, Achi, their target de novo genes, and secondary transcription factors involved in the network. (F) The transcriptional regulatory network of Vis, Achi, Jra, their target de novo genes, and secondary transcription factors involved in the network. (G) Tissue expression pattern of predicted vis target de novo genes in Fly Cell Atlas dataset. (H) Expression pattern of predicted vis target de novo genes in different testis cells in Fly Cell Atlas dataset. In this figure, SPG stands for spermatogonia, SPC for spermatocytes, and SPD for spermatids. The p-values of whether the predicted targets were significantly turned on or off after the expression of *vis* in spermatocytes were shown in the top panel.

### Vis/Achi/Jra may act as major transcription factors for de novo gene candidates

Interestingly, the top three transcription factors, Vis, Jra, and Achi, were predicted to regulate 139, nearly 28% of the 498 expressed de novo gene candidates. We termed the 139 de novo gene candidates as VAJ-regulated de novo gene candidates. When further investigating the regulatory metrics of the 139 VAJ-regulated de novo gene candidates, we found that these genes have significantly fewer regulatory transcription factors (Figure 2C) and alternative transcriptional regulations (defined in Methods) (Figure 2D) than other de novo gene candidates and testis-biased genes, suggesting that many de novo gene candidates exhibit more directed transcriptional regulations.

Interestingly, among the top three transcription factor genes, *vis* (*vismay*) and *achi* (*achintya*) are tandem duplications. In the SCENIC analysis, the top transcription factor, *vis*, was predicted to directly regulate 78 of de novo genes, the largest number among all the de novo gene TFs, while *achi* were responsible for the transcriptional regulation of 45 de novo gene candidates, 21 of which are also regulated by *vis* (Figure 2E). In total, *vis* and *achi* were predicted to directly regulate 102 de novo gene candidates. The direct regulations were also complemented by a few alternative transcriptional regulations by secondary transcription factors (Figure 2E). The core DNA binding motifs of both Vis and Achi were TGACA according to the latest JASPAR database ^16^ and supported by additional AlphaFold3 predictions (Figure 2E, Figure S5).

The other transcription factor in the top three regulators was *Jra*, which was predicted to directly regulate 66 de novo gene candidates, with 29 of them also regulated by *vis* or *achi* (Figure 2F). *Jra* has several important roles in *Drosophila*, including development ^17^ and immune response^18^. Interestingly, its mammalian homologs, *jun*, were reported to be oncogenic ^19^. The core DNA binding motif of the mammalian homologs, TGACTCA, was similar to *vis*/*achi*, TGACA (Figure 2E and 2F). In all, the results altogether indicate a straightforward transcriptional mechanism of around a quarter of the de novo gene candidates identified in *D. melanogaster*, with both minimized transcription factor binding motifs and direct transcriptional regulations.

To partially confirm the transcriptional regulation of *vis*, we analyzed the expression pattern of 78 de novo gene candidates that were predicted to be directly regulated by *vis* in Fly Cell Atlas datasets. We found almost all the regulated de novo genes showed strongly biased expression towards testis (Figure 2G). The expression pattern is expected as *vis* is strongly biased in testis (Figure 2G). We further investigated the expression pattern of *vis* and its predicted 78 de novo gene targets in testis, and found that most (65) of the 78 de novo gene targets were turned on in the spermatocyte stage, with several (5) of them being turned off (Figure 2H, Table S2). This is consistent with the expression pattern of *vis*, starting to be highly expressed in early spermatocytes and peak in late spermatocytes (Figure 2H). These results further indicate that Vis/Achi/Jra could act as major transcription factors that regulate the expression of nearly 28%, 139 out of 498, expressed de novo gene candidates.

### Vis/achi upregulated de novo genes expression

As mentioned above, two tandem duplicated TF genes, *vis* and *achi*, were among the top three identified major transcription factors. Functionally, *vis* and *achi* are meiotic arrest genes and can regulate genes involved in meiotic division and differentiation in spermatogenesis through interacting with other tMAC complex proteins, including Always early (*aly*) and Cookie monster (*comr*) ^20,21^. The proteins encoded by the two genes, Vis and Achi, were highly similar with a protein sequence identity of 96.4%. These genes represent a recent duplication event in the *D. melanogaster* species complex (Figure 3A). In addition, they showed similar but slightly different expression patterns, with both being testis-specific (Figure 3B), while *vis* was expressed at an approximately 1.6-fold higher level in the testis of *D. melanogaster* and *achi* showed a low expression level in tissues outside of the testis. However, the two genes appear to be partially redundant, as each can compensate for the other to some extent ^21^. Notably, many de novo genes are enriched in pre- and post-meiotic germ cells in the testis ^7,12,14^, aligning with the regulatory roles of Vis and Achi. Understanding how spermatocyte-expressed transcription factors impact de novo gene regulation is thus a key question to answer.

**Figure 3.**
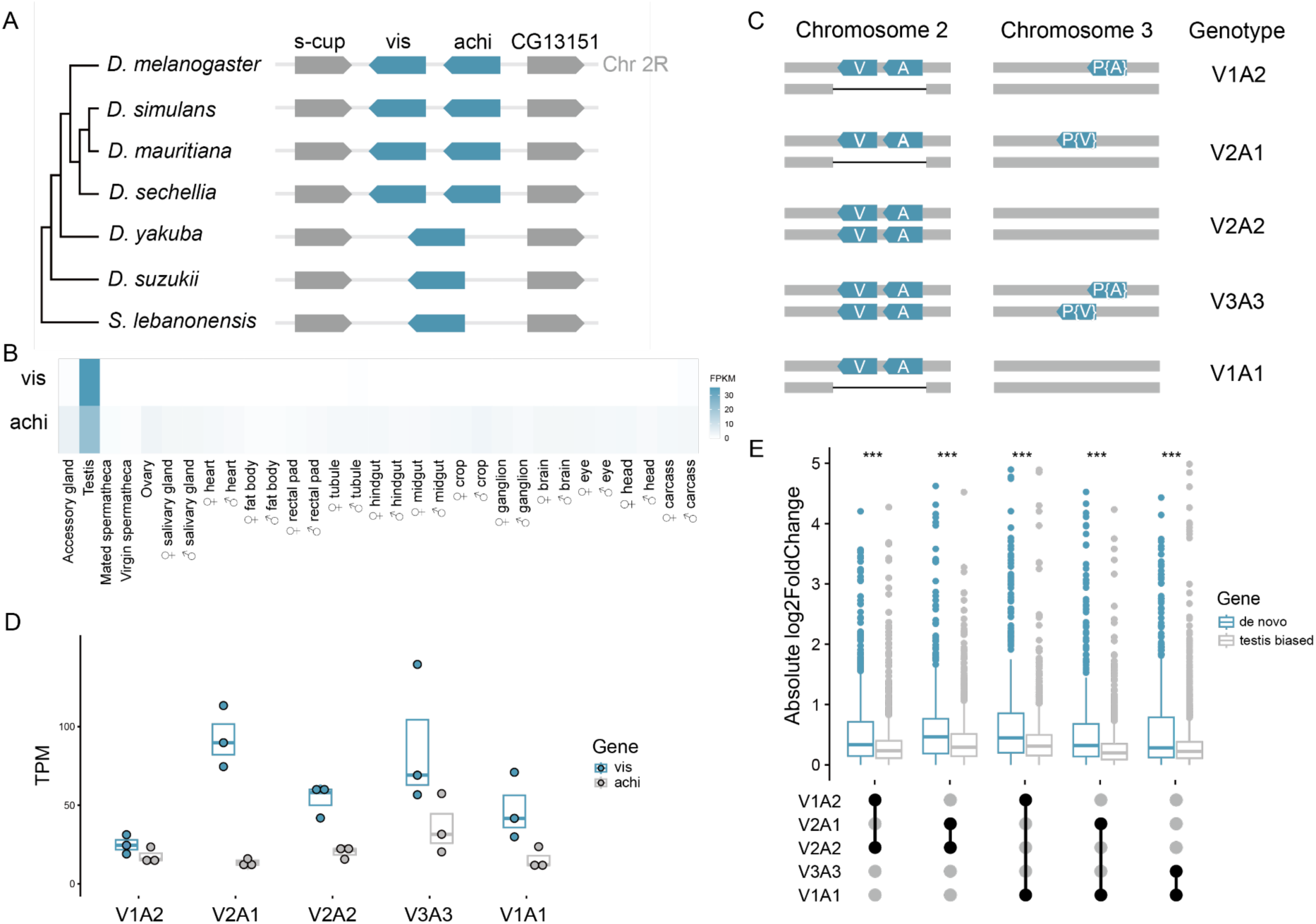
RNA-sequencing of testis from five strains with different copy numbers of *vis* and *achi*. (A) Diagram of synteny map of *vis* and *achi*. (B) Tissue expression pattern of *vis* and *achi* among different tissues in FlyAtlas2. (C) Diagram of five different strains with different copy numbers of *achi* and *vis*. For simplicity, we used AiVj to denote *i* copies of *achi* and *j* copies of *vis*. (D) Expression levels (TPM) of vis and achi in different RNA-seq samples. (E) De novo gene expression showed larger variations (absolute log2foldchange) in strains with more copy numbers of achi or vis when comparing: V2A2 vs. V1A2, V2A2 vs. V2A1, V1A2 vs. V1A1, V2A1 vs. V1A1, and V3A3 vs. V1A1.

To further confirm the transcriptional regulatory functions of *vis* and *achi* in the testis and their effects on the transcription of de novo genes, we manipulated the expression of those two genes and studied the effect on the mRNA abundance of their target genes. We first generated five strains with varying *vis* and *achi* copy numbers while controlling for genetic backgrounds. This was achieved using knockout strains for both *achi* and *vis* on chromosome 2R and an extra copy of *vis* or *achi* on chromosome 3 ^21^ (Figure 3C and Figure S6). For simplicity, we used ViAj to denote *i* copies of *vis* and *j* copies of *achi*. We then performed strand-specific RNA-seq on testes sampled from each strain and compared the gene expression level of de novo genes and other genes between different genotypes. The expression levels of *vis* and *achi* vary with copy numbers across different genotypes (Figure 3D). Among all genotypes, the expression level of vis shows greater variation. The expression is higher with two or three copies of *vis* (V2A1, V2A2, and V3A3) and lower with only one copy (V1A2 and V1A1). The expression level of *achi* is similar across different genotypes, except for higher expression with three *achi* copies (V3A3). Although the expression levels of *vis* and *achi* vary among different genotypes, the samples did not cluster drastically separately by genotype in the PCA plot (Figure S7), suggesting that the changes of *vis* and *achi* expression did not have widespread effects and the overall transcriptome in different genotypes remain relatively homogeneous.

Considering the testis-biased expression of most de novo genes, we compared their expression level changes with those of other testis-biased genes. In all pairwise comparisons between different genotypes, de novo genes exhibited stronger expression variation, with significantly higher absolute fold changes compared to other testis-biased genes (Figure 3E). The fold changes were shown in the comparisons where only copy numbers of *vis* or *achi* differed (Figure 3E), e.g., V2A2 to V1A2, V2A1 to V1A1, V2A2 to V2A1, and V1A2 to V1A1, as well as in the comparison where *vis* and *achi* differed by two copies (V3A3 to V1A1). These comparisons indicated that both *vis* and *achi* can affect the transcription of de novo genes.

We further analyzed the RNAseq data using multiple regressions by applying a linear model to determine if the mRNA abundance of *vis* or *achi* can be a predictor of the expression of their target genes. In order to find the best model across all genes to analyze the data, we performed an Akaike information criterion (AIC) analysis comparing 4 different models: T∼V, T∼A, T∼V+A and T∼V+A+AxV, where T represents the expression of target genes, A for *achi* expression, and V for *vis* expression. We found that for 95% (2745/2887=0.95) of the testis-specific genes, the T∼V model had the lowest AICc (AIC corrected for small sample sizes) scores or was not significantly different from the best-fitted model (i.e., had a delta AICc < 2). This result supports that the T∼V model is the best fit across most testis-specific genes, we thus used the T∼V model to further examine how changes in *vis* expression affect the expression of other genes. In particular, we were interested in whether target genes in the Vis/Achi regulon were more likely to be affected by changes in *vis* expression. Specifically, we tested the hypothesis that genes with non-zero regression slopes in the Vis/Achi regulon exhibit better fit to the model (higher R-square values) than genes not in the Vis/Achi regulon.

We found that testis-specific genes in the Vis/Achi regulon had significantly higher mean R-square values than genes not in the Vis/Achi regulon (Figure 4A: mean±s.e.m. 0.33±0.01 and 0.29±0.01 respectively; Wilcoxon rank test: W = 196820, p-value = 0.035). This trend was the same but even more pronounced for de novo genes: mean±s.e.m. 0.35±0.02 and 0.26±0.01, respectively (Figure 4A: Wilcoxon rank test: W = 1990, p-value = 0.00076). However, we did not find transcription factors to exhibit the same trend (Figure 4A: mean±s.e.m. 0.40±0.06 and 0.34±0.02 respectively; Wilcoxon rank test: W = 541, p-value = 0.33). The observed higher R-square values in the Vis/Achi regulon support our hypothesis that those genes fit the model better than others. This is thus an independent experimental confirmation of the Vis/Achi regulon. Interestingly, de novo genes appear to respond even more linearly to Vis mRNA abundance suggesting that their regulation is more direct or more simple, while transcription factors - probably the genes with the more complex regulatory network connections- do not follow this pattern. While one can expect de novo genes to respond linearly, transcription factors may respond in a non-linear manner. These results align with previous work which found that younger genes and genes with fewer network connections tend to respond more linearly to TF expression ^22^.

**Figure 4.**
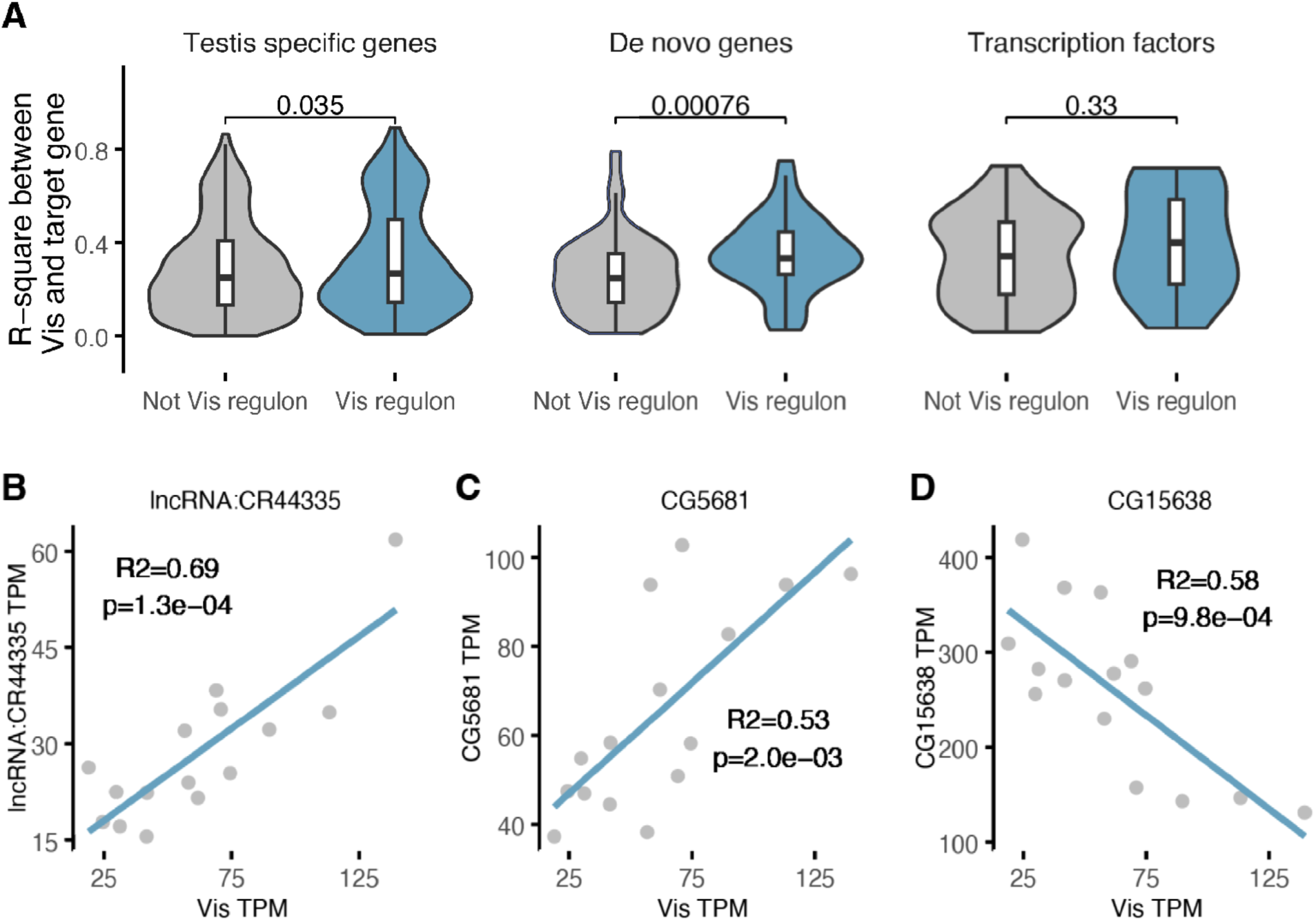
Expression of the genes in the Achi-Vis regulon is correlated with Vis’ mRNA abundance. A) the violin plots show distribution of R-square from the regression between Vis TPM and the gene target’s TPM (outliers are not displayed for visual clarity). P values are from the Wilcoxon ranks test. B) Regression of de novo gene (FBgn0265423) CR44335’s TPM over Vis TPM. C) Regression of de novo gene (FBgn0032658) CG5681’s TPM over Vis TPM. D) Regression of de novo gene (FBgn0028943) CG15638’s TPM over Vis TPM.

In order to identify the genes with large and potentially most meaningful regressions, we applied a 5% FDR cutoff on the adjusted p-values of the regression. For example, we found that *lncRNA:CR44335* which is presumed to have been born in the branch leading to *D. pseudoobscura* group ^14^ is in the Vis-Achi regulon and harbors a significant regression to *vis*’ expression (Figure 4B). Similarly, *CG5681* is presumed to have originated in the phylogenetic branch leading to *D. ananassae* ^14^, is in the Vis-Achi regulon and exhibits a significant regression (Figure 4C). We found that 97% (1091/1130=0.965) of all the genes passing the FDR cutoff exhibit a positive slope, meaning that the TPM of the target gene increases when *vis*’ mRNA abundance increases (Figure S8). This result is thus consistent with *vis* being a transcription factor that plays the role of transcription activator in the testis (Figure 2H).

However, previous work on transcription factor expression coupling suggests it is not uncommon even for suppressors of transcription to exhibit positive correlations with their targets^22^. Similarly, we found that *CG15638* born in the branch leading to *D. willistoni* which belongs to Vis-Achi’s regulon exhibits a significantly negative correlation with Vis’ expression in our RNAseq experiment (Figure 4D). However, in single-cell RNA-seq data, *CG15638*’s expression is turned on at the spermatogenesis stage where Vis’ expression is turned on ^8^.

It was reported that Vis/achi proteins can interact with testis meiotic arrest complex, tMAC (^21^). To investigate whether the expression of different subunits within the tMAC complex could be affected by *vis*/*achi* and potentially contribute to the regulation of de novo genes, we analyzed the correlations between the expression levels of *vis* and genes encoding tMAC subunits, *aly*, *come*, *tomb*, *p55*, *wuc*, *topi*, and *mip40*. The former five subunit genes, *aly*, *come*, *tomb*, *p55*, and *wuc*, showed significant positive correlations, while the latter two subunit genes, *topi* and *mip40*, did not (Table S3). The correlations are consistent with previous studies showing that the function of the latter two subunits, topi and Mip40, may be independent of Vis/Achi. For example, it was reported that the *topi* subunit may have parallel function with Vis/Achi for recruiting Aly and other subunits in the complex to chromatin ^23^, and Mip40 may belong to a variant of tMAC that lacks Vis/Achi ^24^. In addition to the five significantly correlated tMAC subunit genes, the expression levels of several transcription factor genes functioning in spermatocytes showed significant correlations with *vis*/*achi* (Table S3), including *can* and *sa* in testis-specific TATA-binding protein associated factors (tTAF) complex ^25^, all four subunits in testis-specific polymerase-associated factor (tPAF) complex, Paf1L (*CG12674*), Cdc73L (*CG6220*), Ctr9L (*CG9899*), and Leo1L (*CG10887*) ^26^, and two testis-specific bromodomain proteins, tBRD-1 and tBRD-2 ^27^. These correlations suggested that there were possible cross-talks between these transcription factors and that several other transcription factors may also be involved in the regulation of de novo gene expression. In contrast, the expression levels of transcription factor genes functioning in other processes during spermatogenesis did not show significant correlations with *vis* (Table S3), for example, *bam* in spermatogonia, *STAT* in germline stem cells, and *chinmo* in somatic cyst stem cells ^28–30^.

### Vis/Achi may contribute to the emergence of young de novo genes in *D. melanogaster*

We have shown that de novo genes that were predicted to be regulated by the top three major transcription factors have relatively straightforward transcriptional regulations (Figure 2C and 2D), which we termed as VAJ-regulated de novo gene candidates. We next ask whether these straightforward transcriptional regulations would link to de novo gene emergence. We compared the origination age of the VAJ-regulated de novo gene candidates and other de novo gene candidates inferred from an earlier study, with Br1 being the youngest, which were mel-complex specific, and Br9 the oldest ^14^. We did not observe significant difference between the origination ages among these de novo genes (one sided Wilcoxon rank test p = 0.88, Figure 5A). However, we found that young de novo genes in Br1 were more likely to be regulated by the top three transcription factors (one-tail fisher exact test p = 0.04, Figure 5A). Notably, seven out of the eight de novo genes that were predicted to originate from Br1 are among the VAJ-regulated de novo gene candidates (Figure 5A). We further found that young VAJ-regulated de novo genes in Br1 showed even more straightforward transcriptional regulation than older VAJ-regulated de novo genes with less direct regulating transcription factors (Figure 5B) and less transcriptional regulation paths (Figure 5C). Among the seven young VAJ-regulated de novo genes in Br1 that showed expression in fly cell atlas data, five of them were regulated by Vis, Achi, or both (Figure 5D). The results indicate that Vis/Achi may contribute to the emergence of young de novo genes in *D. melanogaster*.

**Figure 5.**
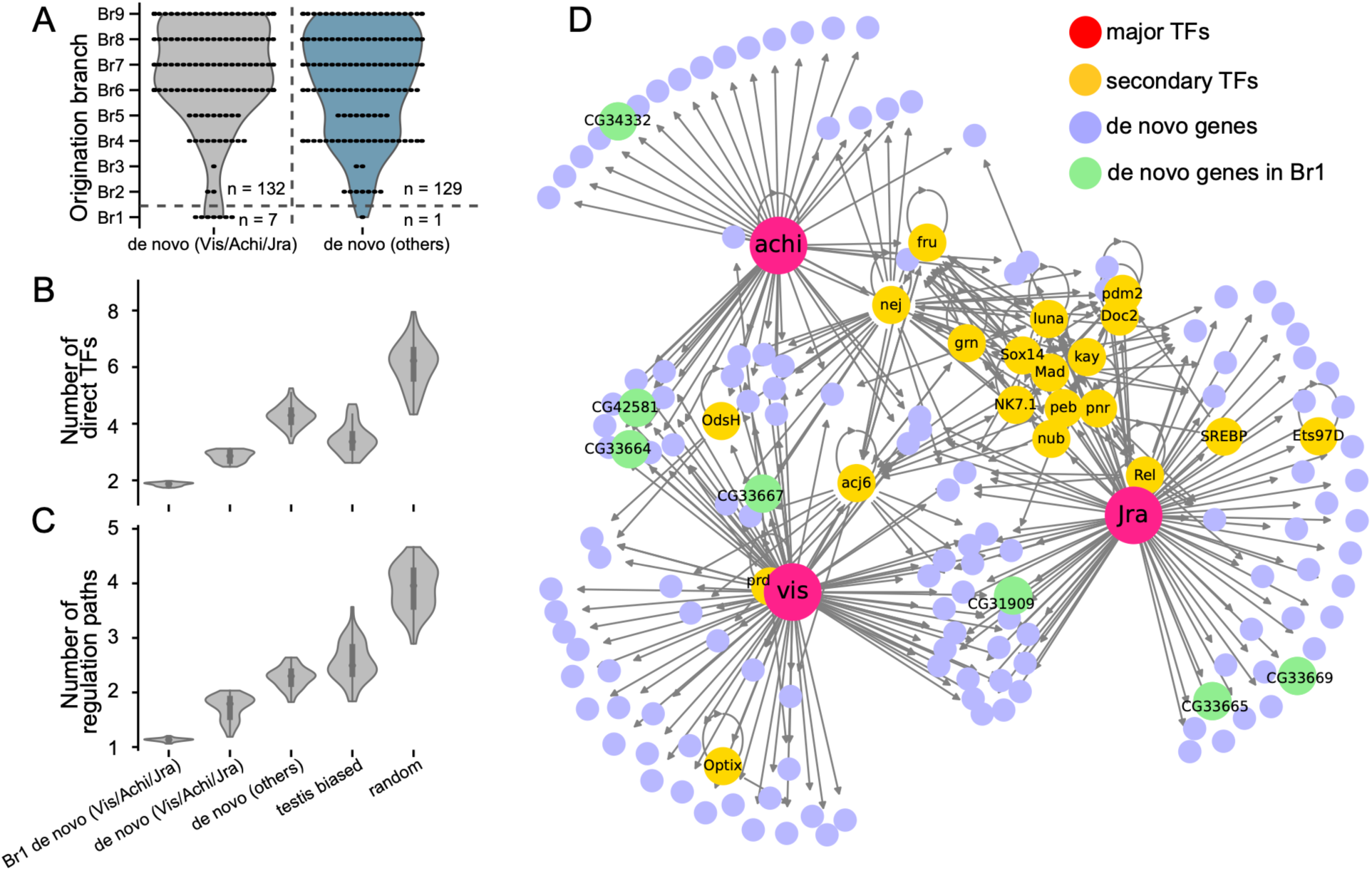
Young de novo genes originated in mel-complex species were more likely to be regulated by Vis/Achi. (A) The origination branch of VAJ-regulated de novo genes (de novo genes regulated by top three transcription factors, Vis/Achi/Jra) and other de novo genes. The two categories did not show significant differences (one sided Wilcoxon rank test p = 0.88). When focused specifically on Br1 and other branches, young de novo genes originated in Br1 were more likely to be VAJ-regulated de novo genes (one-tail fisher exact test p = 0.04). The young VAJ-regulated de novo genes originated in Br1, which we denoted as Br1 de novo (Vis/Achi/Jra), showed less complex transcriptional regulations compared to other de novo genes, with less number of direct TFs (D) and regulation paths (C). Among the seven VAJ-regulated de novo genes that originated in Br1, five were predicted to be regulated by Vis/Achi (D).

## Discussion

Multiple models have been proposed for de novo gene emergence, such as the transcription-first ^31^, ORF-first ^32^, and the cultivator model ^33^. These models are not mutually exclusive and often have synergistic effects in various scenarios. However, the recruitment of transcription and transcriptional regulation is essential regardless of the specific model, and it remains unclear whether specific sets of transcription factors are involved. In this study, we made use of the recent fly cell atlas data ^13^ and employed SCENIC ^34,35^ analysis to investigate the basis of transcriptional regulations of de novo genes in *D. melanogaster*. From the SCENIC analysis, we showed that most of the de novo gene candidates were predicted to be regulated by only a few transcription factors. Among these transcription factors, Vis, Achi, and Jra had the largest number of predicted de novo gene targets, accounting for more than a quarter of the de novo gene candidates (Figure 2A). The results suggest that the transcription factors initiating the transcription of de novo genes are not completely random, at least in *D. melanogaster*. Whether this lack of randomness is primarily driven by how natural selection maintains novel genes or by the greater flexibility of these transcription factors in binding new sites remains unknown.

However, Achi/Vis regulate many genes important for meiosis in the male germline, and it is reasonable to expect that novel genes potentially affecting meiosis and post-meiotic processes would have a higher probability of being either selected for or selected against by natural selection. In another word, the maintenance of the regulatory machinery of de novo genes may be due to the important functions of de novo genes, rather than the regulatory ability of Achi/Vis.

We further conducted a network analysis of the transcriptional regulatory network and found that de novo genes regulated by Vis, Achi, and Jra, which we denoted as VAJ-regulated de novo genes, show lower regulatory complexity compared to other targets. This observation is in line with the widely accepted hypothesis that de novo genes are more likely to start with “simple beginnings” ^36^. If a gene exhibits a low level of regulatory complexity, its promoter and enhancer binding sites are more likely to be easily acquired through mutation or even exist without mutations. For example, TGACA, which is the binding site of Vis, Achi, and part of Jra, is highly prevalent across the genome. New enhancers or promoters with these binding sites may arise through a single mutation or even without any mutation, facilitated by three-dimensional chromosomal reorganization. Another contributing factor may lie in the intrinsic regulatory properties of these transcription factors. Vis and Achi, for instance, possess the capacity to regulate a large number of genes, providing the flexibility to expand their regulated gene sets.

Our results suggest that both of these “simple” properties—low regulatory complexity and versatile regulatory capacity—may have facilitated the regulation of de novo genes by Vis, Achi, and Jra.

It is interesting to hypothesize how a simple promoter and enhancer can originate. Previous studies, such as those conducted in *E. coli* ^37^ and *Drosophila* ^38^, have shown that novel promoters and enhancers can arise from a few mutations within random sequences. In *Drosophila*, a few key nucleotide changes may be sufficient to alter the chromatin state—for example, transitioning from closed chromatin to open chromatin ^39^. These findings demonstrate a significant potential for existing or even random sequences to be repurposed into new regulatory elements. Strikingly, in *Drosophila* testis germ cells, during a stage closely associated with the expression and translation of Achi and Vis, at least one-third of genes exhibit alternative promoters during the transition from spermatogonia to spermatocytes ^40^.

Many of the spermatocyte-specific promoters contain tMAC and Achi/Vis motifs, further highlighting the versatile regulatory capacity of these TFs. In addition to a high regulatory capacity of tMAC complex, potentially including Achi and Vis, male germline also has a remarkable promoter diversity for genes.

To verify the transcriptional regulation of Vis/Achi, we constructed five different strains of *D. melanogaster* with different copy numbers of *vis* and *achi* genes. We showed that *vis* and *achi* could upregulate de novo gene expression (Figure 3E), especially the de novo genes that were predicted to be regulated by them (Figure 4). This result is consistent with single-cell data analysis where we found that most of *vis*-regulated de novo genes were turned on after Vis expression in spermatocytes (Figure 2H). Our results suggest that de novo genes that are predicted to be regulated by Vis tend to respond in a simple, linear model, while transcription factors regulated by Vis respond in a more complex manner (Figure 4, Figure S8), suggesting a straightforward regulatory pathway between Vis and its predicted de novo gene targets. For instance, it is possible that de novo genes are primarily regulated by a single transcription factor complex, resulting in the linear expression correlation observed. Indeed, this simplicity may be explained by the smaller number of directly regulated transcription factors and the fewer transcriptional regulation paths that we observed in this study (Figure 2C and 2D).

Although there is substantial overlap in the predicted target de novo genes between Vis and Achi, we also found some predicted to be Vis-specific and others Achi-specific. Due to the very high protein sequence identity between Vis and Achi and the similarities in their binding motifs, we were unable to fully determine whether the observed differences in genes regulated by Vis and Achi identified through regulon analysis are of biological significance or a computational artifact. Subtle differences in their sequences as well their expression differences (Figure 3D) could contribute to different regulation patterns in the regulon analysis. However, we cannot exclude the possibility of randomness in single-cell regulon analysis when assigning gene targets to Vis or Achi regulons. Therefore, in many analyses, the two regulons were combined and analyzed as a whole. In addition, due to the limitation of currently available cis regulatory motifs, many other transcription factors, such as Topi in tMAC, were not included in the current study. To overcome these limitations, further large-scale sequencing or computational studies on transcription factors and enhancers are needed. One possible direction would be to incorporate single cell ATAC-seq data and apply SCENIC+ ^41^ and build cell specific enhancer driven gene regulatory networks.

Although it is theoretically impossible to distinguish parental from new copy for tandem duplicated genes, several pieces of evidence support *achi* could be the de facto ancestral copy (the slower evolving copy). First, its broader expression pattern (Figure 3B), compared to the tendency of new copies to have more specific expression patterns, and second, its shorter branch length in the gene tree (Figure S9), support this hypothesis. Third, after knocking out *Achi/Vis*, *achi* is more effective at rescuing the mutant phenotypes than *vis* ^21^. Thus, it is likely that Achi has retained more of its ancestral function, while Vis has undergone subfunctionalization and/or neofunctionalization. Notably, knocking out *vis* resulted in a significant upregulation of *achi* in the *vis* mutant ^42^, suggesting the presence of a feedback loop to compensate for or stabilize the transcription and translation of the two genes. Consequently, the differences observed in the genetic experiments likely reflect only a subset of the changes regulated by Achi/Vis. Nevertheless, it is evident that Achi and Vis are critical regulators of spermatocyte-expressed genes, including *de novo* originated genes (^20,21^).

From our analysis, we found that Vis and Achi may be key transcription factors that contribute to the emergence of young de novo genes in *D. melanogaster*. Specifically, we found that most of the young, mel-complex species specific de novo genes were predicted to be regulated by Vis/Achi. Notably, *vis* and *achi* genes are tandem duplicates specifically found in *D. melanogaster* species complex. Cases of transcription factor duplications where one paralog retains testis expression and function are in fact not that uncommon ^26,43^. This raises the possibility that such transcription duplication events could possibly be related to the emergence or the maintenance of those young de novo genes. In the gene tree (Figure S9), the branch lengths in the *simulans* species complex are longer, indicating higher sequence divergence compared to *D. melanogaster*. Moreover, *achi* and *vis* cluster by gene rather than by species, suggesting that the majority of sequence changes occurred before the speciation events of *D. simulans*, *D. sechellia*, and *D. mauritiana* (Figure S9). Interestingly, the clustering of *achi/vis* in *D. melanogaster* as an outgroup implies a greater degree of functional divergence in *achi* and *vis* within the *simulans* species complex. This observation warrants further investigation. Note that, in other species, there may be other transcriptional regulation mechanisms for de novo genes and their emergence. Our study provides a workable solution to uncover the transcriptional regulation of new genes.

## Material and Methods

### Analysis of fly cell atlas

The snRNA-seq data for *D. melanogaster*, “10x VSN All, Relaxed”, were downloaded from Fly Cell Atlas ^13^ (https://cloud.flycellatlas.org/index.php/s/sgYtFKWx8w3GX22). This annotated dataset contains the UMI count matrix for 13,968 protein-coding genes in 570,000 nuclei across 15 tissues and 17 major cell types. To quantify the expression preference of each gene in different tissues or major cell types, we normalized and scaled the UMI counts as follows. We first calculated the total number of UMI counts in each tissue or major cell type. We then normalized the expression of each gene in each tissue or major cell type by the total UMI counts in the tissue or major cell type.

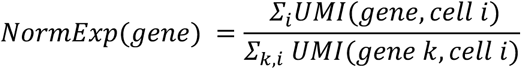

With the normalized expression, the scaled expression of each gene for a specific tissue or major cell type is then given by Zscore of each gene as follows

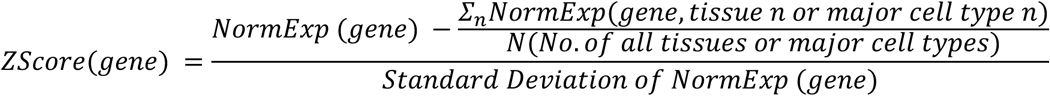

In the analysis, we define a gene as biased expressed in a tissue or a major cell type if its Zscore is greater than 2.5.

### Regulon analysis

We used the python version of SCENIC ^34^, pySCENIC ^35^, to predict the regulons for the de novo gene candidates inferred from our previous study ^14^. A list of transcription factors in *D. melanogaster* were obtained at https://resources.aertslab.org/cistarget/tf_lists/allTFs_dmel.txt. To increase computational efficiency, we constructed a dedicated UMI count matrix that includes only the counts of transcription factors, de novo gene candidates, and testis-biased genes in each cell. The testis-biased genes have Z-score greater than 2.5 and maximum expression in the testis in FlyAtlas2 ^15^. We further filtered out cells where no de novo genes were expressed and transcription factors were lowly expressed. Specifically, we removed cells from the matrix if the maximum UMI count of the top expressed de novo gene was less than 5 and the maximum UMI count of the top expressed TF gene was less than 20. After this step, the filtered UMI count matrix contained 121,767 cells with 1,418 genes including de novo gene candidates, transcription factors, and other testis-biased genes, with considerable tissue cell type heterogeneity (Figure S10). Using this filtered UMI count matrix, we ran 100 SCENIC predictions to account for the stochasticity in the process ^13^. To ensure consistency, we only chose the frequently predicted regulons that appeared in more than 80 of the predictions for the next step analysis. For each frequently predicted regulon, we included the gene targets that appeared in more than five of the predictions.

### Network analysis

Based on the SCENIC prediction, we constructed a directed network by connecting each of the transcription factors and their predicted target genes. We defined two metrics to characterize the complexity of transcriptional regulations of a specific target gene, i.e., the number of direct transcription factors and the number of alternative transcriptional regulations. The number of direct TFs describes the number of transcription factors that were predicted to directly regulate the specific gene. The number of alternative transcriptional regulations describes how many alternative regulations were present in the network. Practically, to get the number of alternative transcriptional regulations, we counted the number of distinct regulation paths from a transcription factor to a target gene. In a regulation path, the specific gene is regulated directly by a transcription factor or indirectly through the regulation of a secondary transcription factor. A simplified diagram is shown in Figure S11.

Here, we compared three groups of genes: 1) the de novo genes that were predicted to be directly regulated by the top three transcription factors (Vis/Achi/Jra), which we termed as VAJ- regulated de novo, or de novo genes (Vis/Achi/Jra), 2) the de novo genes that were predicted to be regulated by the remaining transcription factors, and 3) testis biased genes. To compare the two metrics for each gene group, we implemented a bootstrapped analysis, where we randomly sampled 100 genes from each group and calculated the mean numbers of direct TFs and mean numbers of alternative transcriptional regulations. We repeated this step 100 times to minimize randomness, similar to a standard bootstrap analysis.

### Genetic crosses for generating different copies of achi and vis

Strains *Drosophila* stocks #8509 (genotype: w[*]; Df(2R)pingpong, achi[pingpong]/CyO; P{w[+mW.hs]=vis}3/MKRS) and #8511(genotype: y[1] w[*]/Dp(1;Y)y[+]; Df(2R)pingpong, achi[pingpong]/CyO; P{w[+mW.hs]=achi}3) were obtained from Bloomington Drosophila Stock Center. We then generated a set of 5 genotypes carrying different copy numbers of *Achi* and *Vis* through crosses using the *w*^1118^ strain and a double balancer CyO-TM6B strain (Figure S6). These flies were used for RNA sequencing and subsequent analysis to investigate a possible dosage effect of those transcription factors on the expression of downstream genes, including de novo genes.

### Sample preparation for RNA-sequencing

For each of the 5 genotypes, we selected 4-5 days old males, dissected their testes in cold PBS and placed them in Trizol. We prepared three biological replicates for each of the 5 genotypes (for a total of 15 samples). Each sample is made from 12 pairs of testes, except for one for which we have only 6 pairs of testes (A1V1). Total RNA was extracted using the standard Trizol protocol. We recovered between 600 ng and 1920 ng of RNA per sample. The RNA quality was assessed on a Qubit platform and RNA quantity was measured using a NanodropOne instrument. The RNA was then made into RNAseq libraries and sequenced using NovaSeq X Plus platform by Novogene (Sacramento, CA, USA).

### RNA-seq analysis

We trimmed adaptors from RNA-seq reads using Trimmomatic V0.39 ^44^. In addition, low-quality bases were trimmed using Trimmomatic settings LEADING:3 TRAILING:3 SLIDINGWINDOW:10:25 MINLEN:25. The cleaned reads were then aligned to the *D. melanogaste*r reference genome r6.36 using HISAT2 ^45^. The expression levels of transcripts and genes (transcripts per million, TPM) were quantified using StringTie, and reads that fell within gene regions were counted using featureCounts ^46^. Differential expression analyses were carried out using DEseq2 ^47^.

We analyzed the gene expression using R software as follows: first, all genes that showed TPM=0 in more than 3 samples over the 15 samples were removed from further analysis. We then constructed 4 linear models where the expression of a focal target gene is explained by Vis’ expression, Achi’s expression, or a combination of the two: T∼V; T∼A; T∼V+A and T∼V+A+VxA where T is the TPM for each target gene, V is the TPM of *vis* gene, A is the TPM of *achi* gene, and VxA represents the interaction between Vis and Achi. We then compared the Akaike information criterion corrected for small sample size (AICc) of each model for all genes. We then used the best-fitted model to calculate the R-square values of the model for each gene. We then compared the distribution of R-square values between genes within or outside the Vis-Achi regulon using a Wilcoxon test. A higher mean R-square in one group is interpreted as a sign that the model is a better fit, in other words, that *vis* and/or *achi* are better predictors of target gene expression. We then selected de novo gene candidates with the best regressions by applying a 5% FDR cutoff on the regressions’ p-values.

## Data availability

RNA-sequencing data are submitted to NCBI BioProject under the accession number PRJNA1197503.

## Code availability

Code and scripts for this study are available at https://github.com/LiZhaoLab/DeNovoGene_Transcriptional_Regulation.

## Acknowledgements

We thank members of Zhao lab for helpful discussions. Stocks obtained from the Bloomington Drosophila Stock Center (NIH P40OD018537) were used in this study.

## Author contribution

J.P., L.Z., and N.S. conceived the project. J.P. designed and performed large-scale scRNA-seq and regulon analysis, N.S. designed the genetic cross scheme, B.J.W. and N.S. carried out the genetic experiments and RNA-seq analysis, and J.P., L.Z., N.S., and B.J.W. wrote the paper.

