## Supplemental figures and tables for "Gene regulatory networks and essential transcription factors for de novo originated genes"

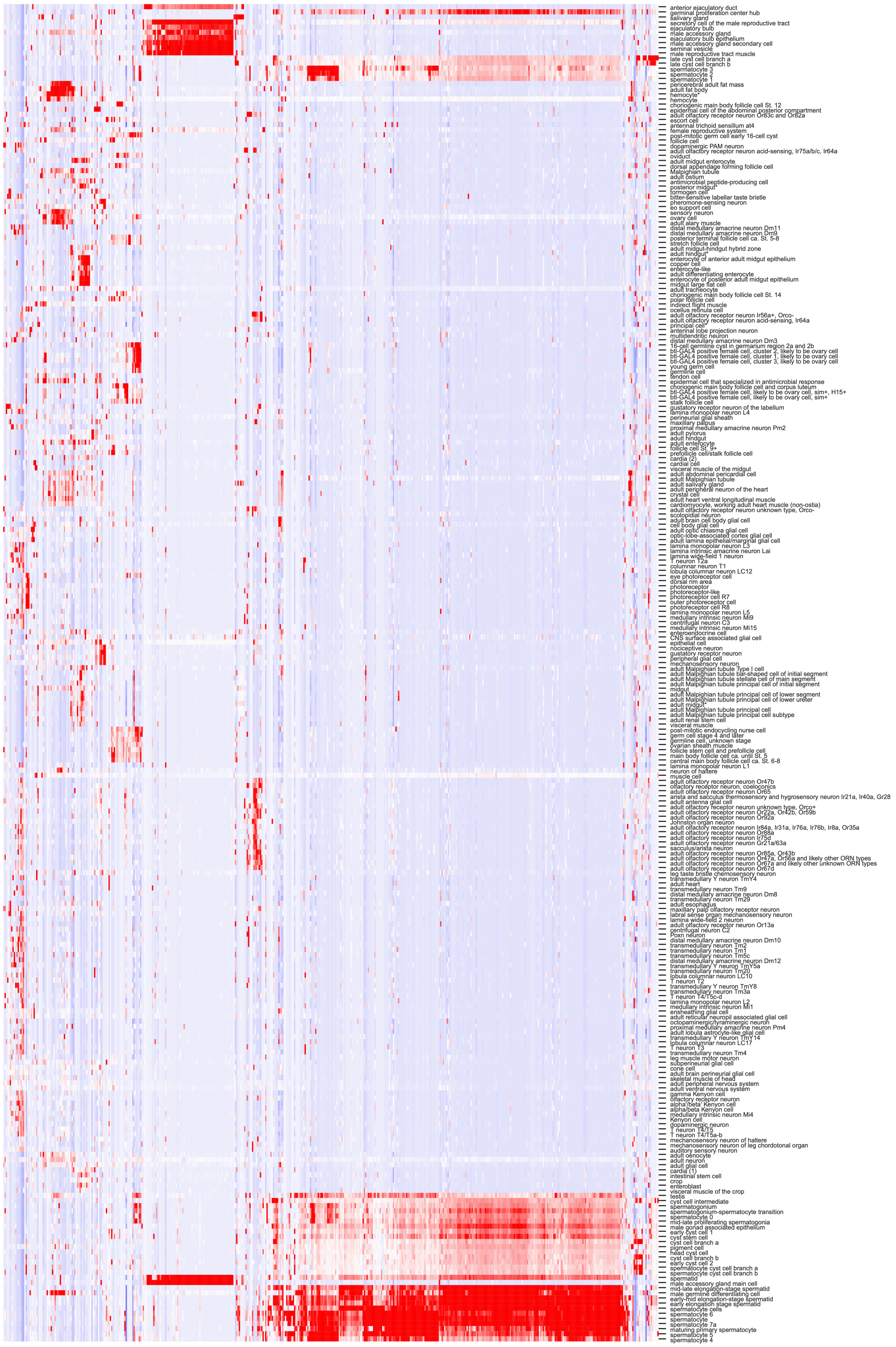


Figure S1. Clustering analysis of de novo gene expression across 250 different cell types. The expression shows clusters in spermatogenesis, gland cells, sensory neurons, hemocytes, ovary cells, midguts, etc.

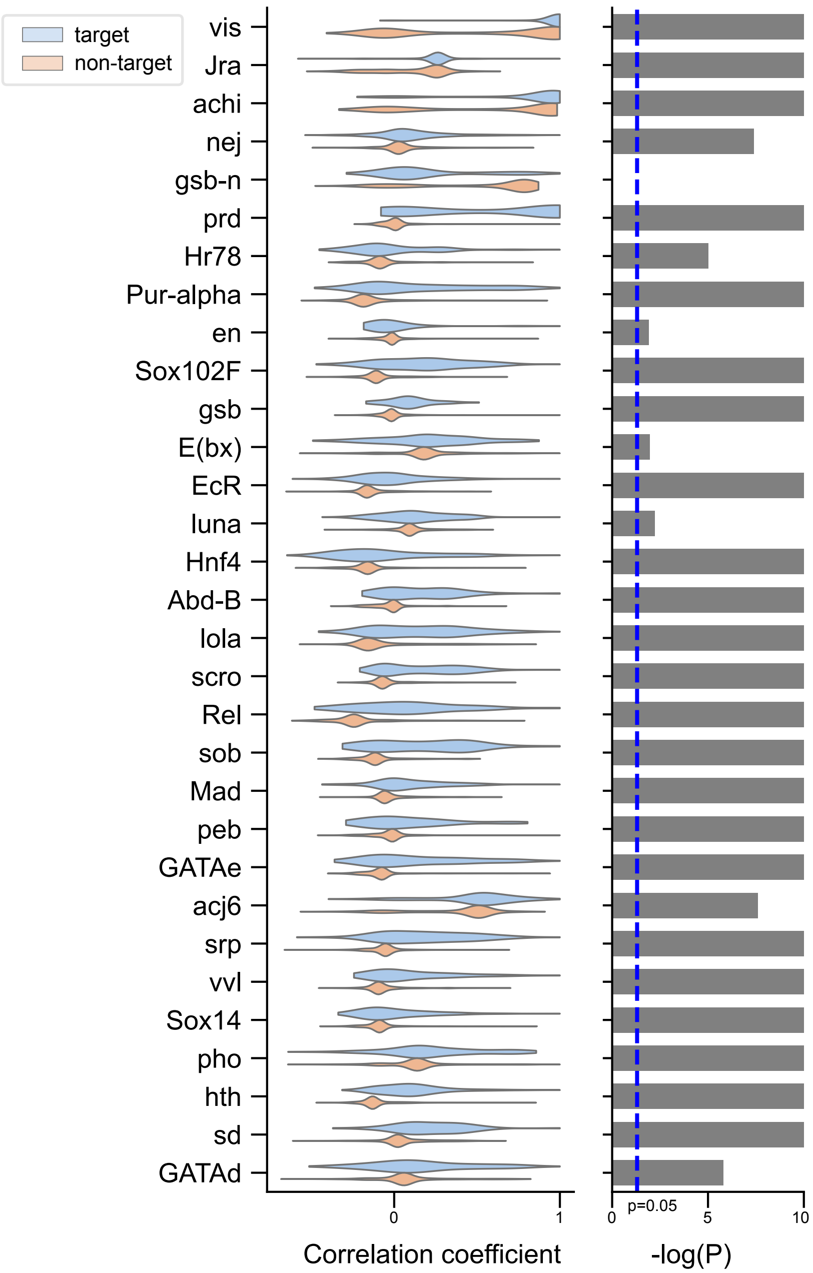


Figure S2. Correlation coefficient of the tissue expression patterns between top 31 transcription factors and their predicted targets. The tissue expression patterns were obtained from FlyAtlas 2. In general, TFs show significant higher correlations with their target genes than other non-target genes. P-values in the right panel were obtained from one sided t-test.

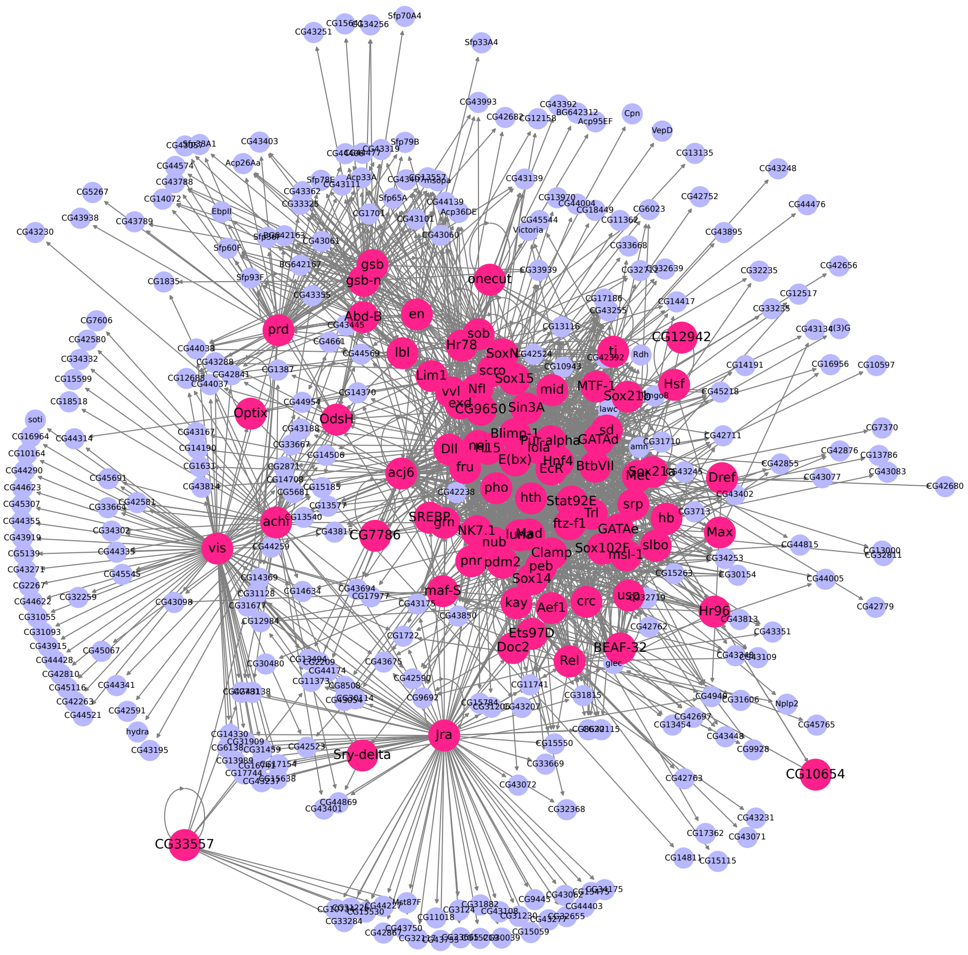


Figure S3. The transcriptional regulatory network between the 81 transcription factors (magenta) and de novo gene candidates (blue).

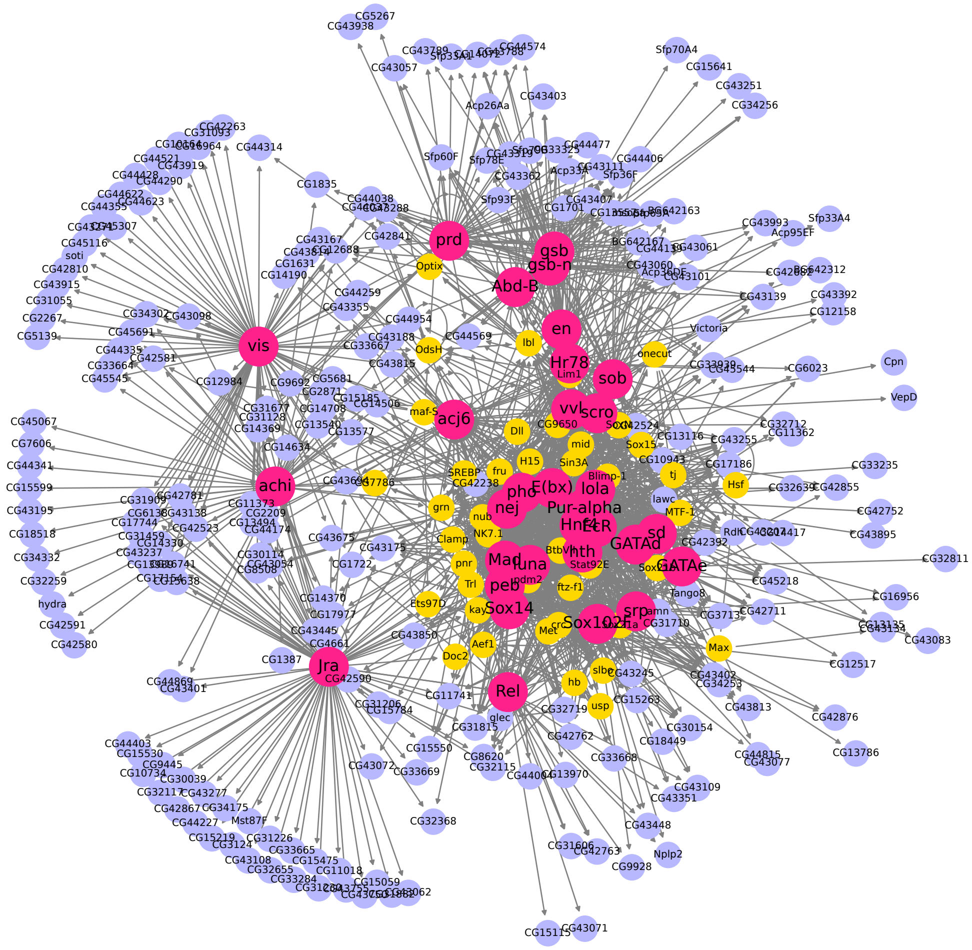


Figure S4. The transcriptional regulatory network between the top 31 transcription factors (magenta) and de novo gene candidates (blue). Other secondary transcription factors (yellow) among the 81 predicted transcription factors were also shown.

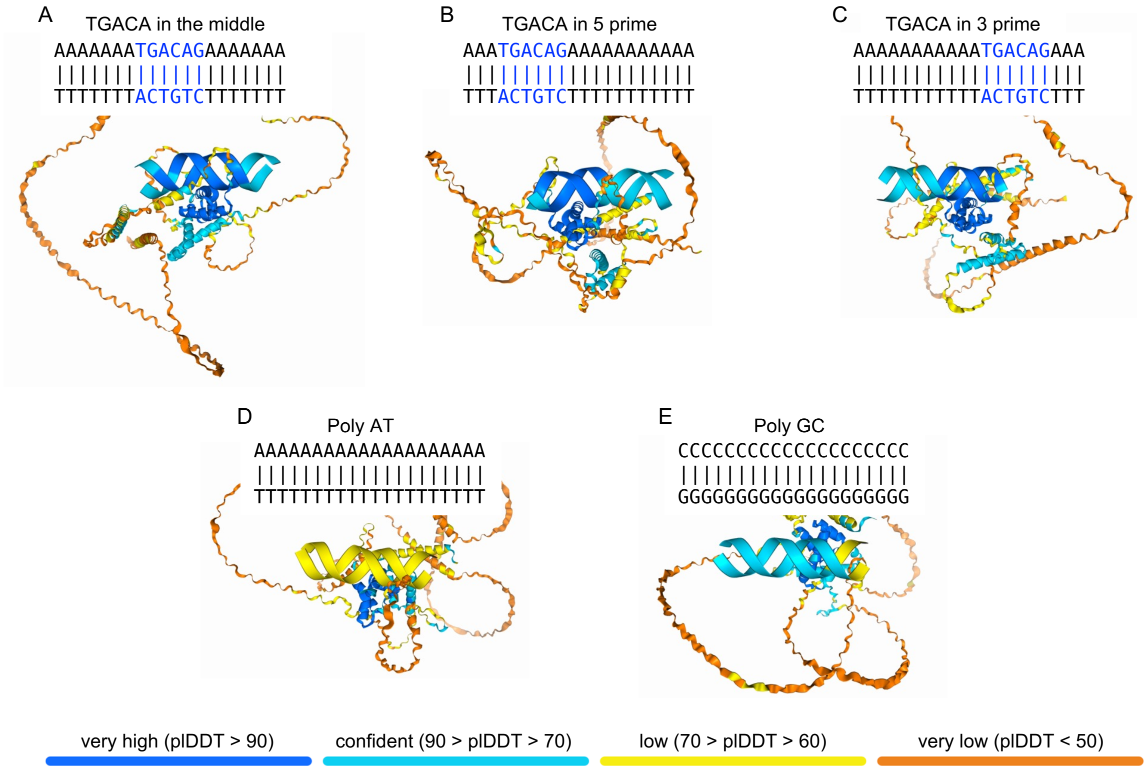


Figure S5. AlphaFold3 prediction of Vis-DNA binding with core DNA binding embedded in the middle (A), the 5-prime (B), and the 3-prime (C) of a poly AT sequence. The prediction between Vis and poly AT (D) and poly GC (E) DNA sequences. The predictions indicate that Vis recognizes the core TGACA motif.

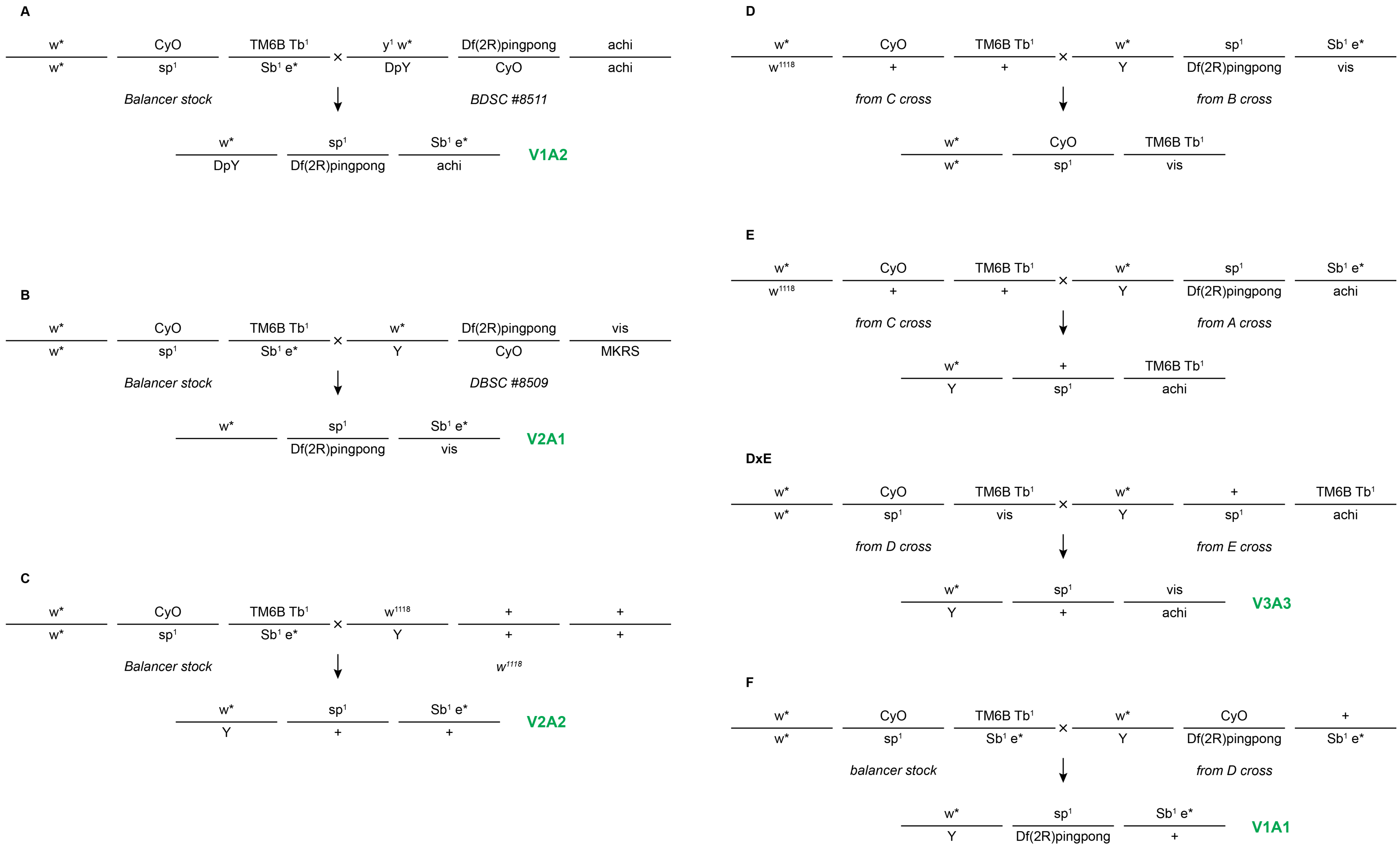


Figure S6. Crossing schemes used to design different copy numbers of *vis* and *achi* with the same genetic background.

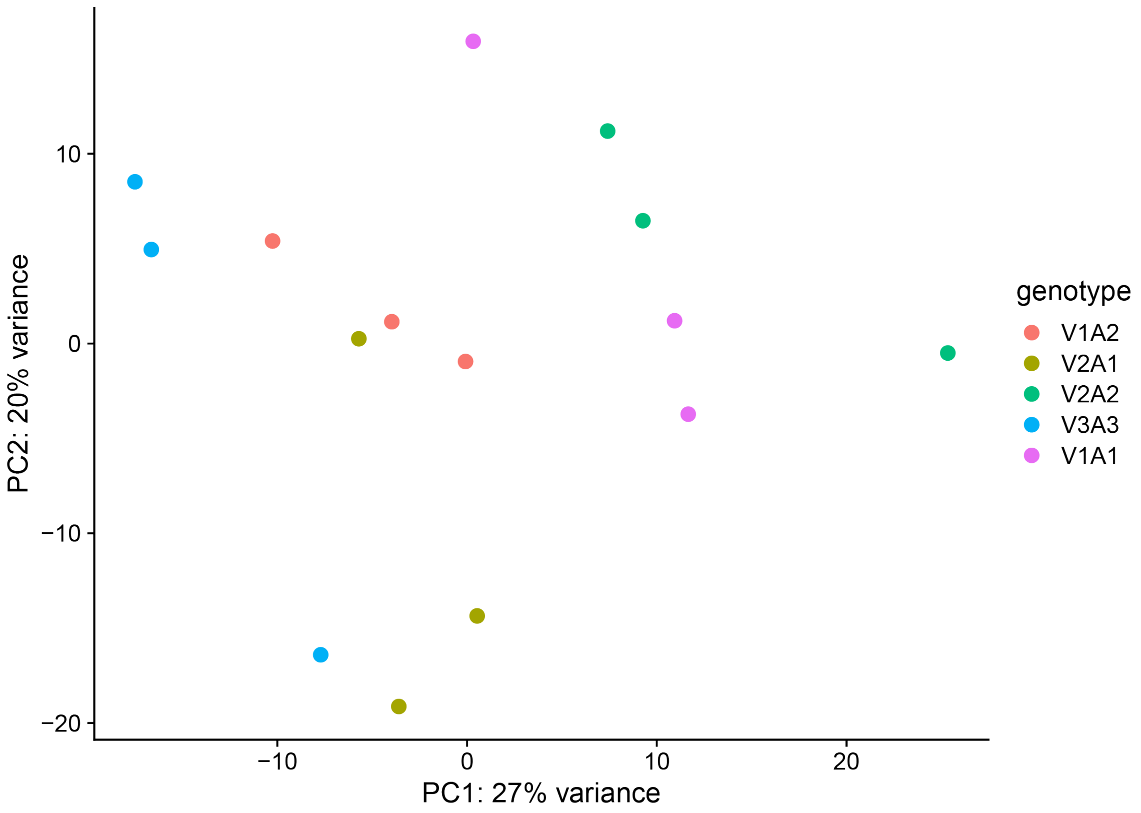


Figure S7. PCA plot of RNA-sequencing data from different strains. The strain information for different labels is the same as in the above Figure (Figure S6). The PCA plot suggest that there were no major changes in overall expression among different strains.

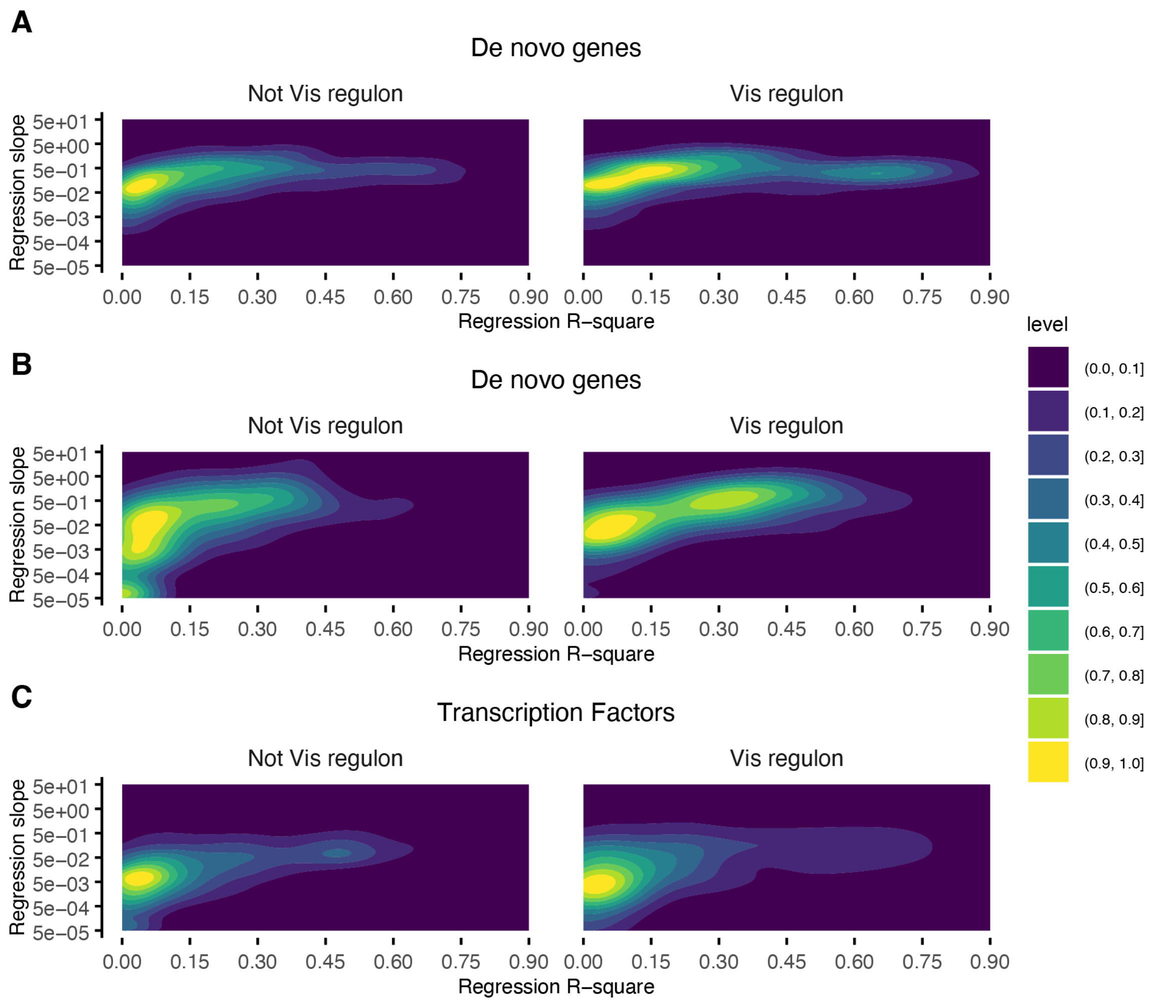


Figure S8. The slope and the coefficient of determination, R-square, between *vis* and *vis* targets, including (A) testis specific genes, (B), de novo genes, and (C) transcription factors, among different RNA-seq samples.

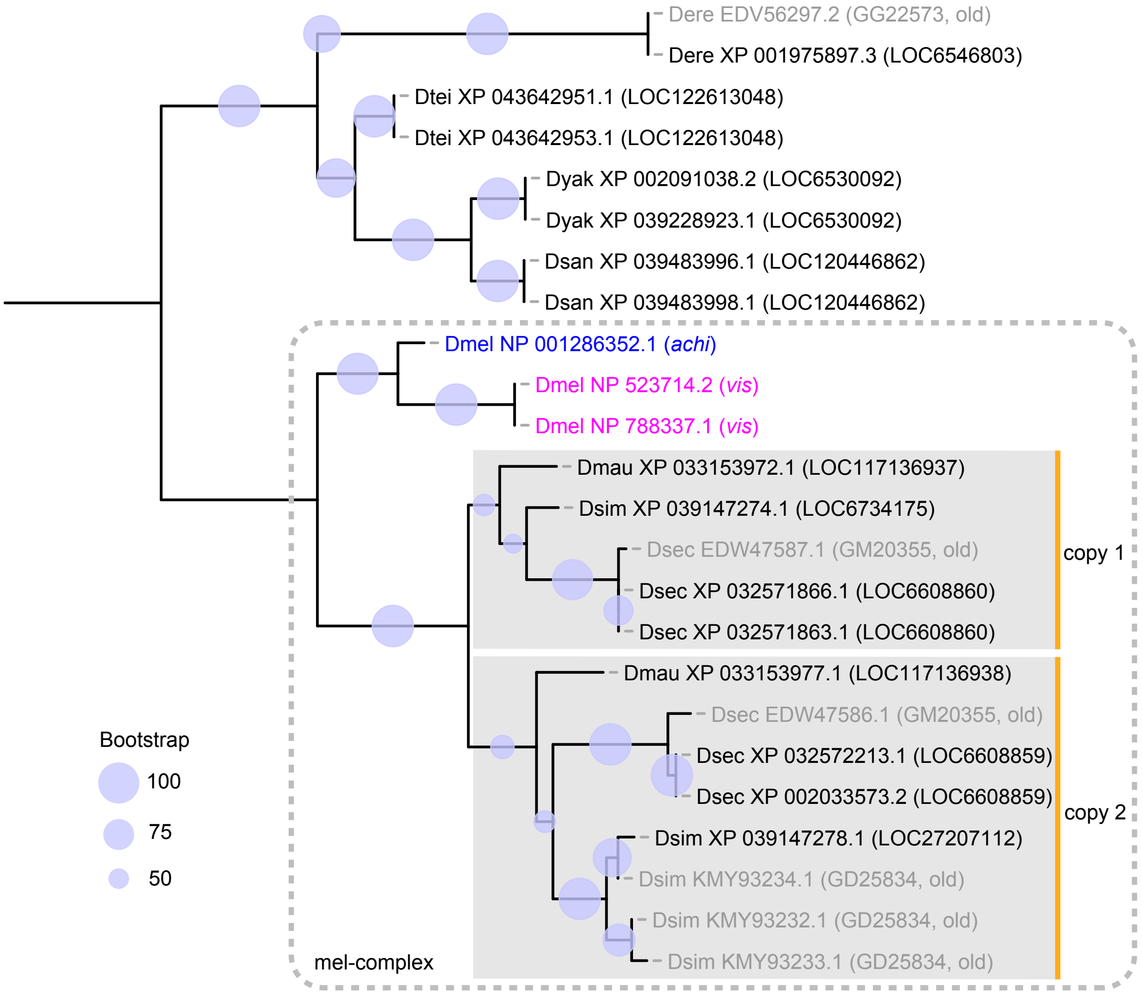


Figure S9. Gene tree of *vis*/*achi* inferred from protein sequence alignments in melanogaster subgroup species. The protein sequences were obtained from blastp search against non-redundant protein sequence database. Protein sequences from suppressed genome assemblies and annotations were shown in gray.

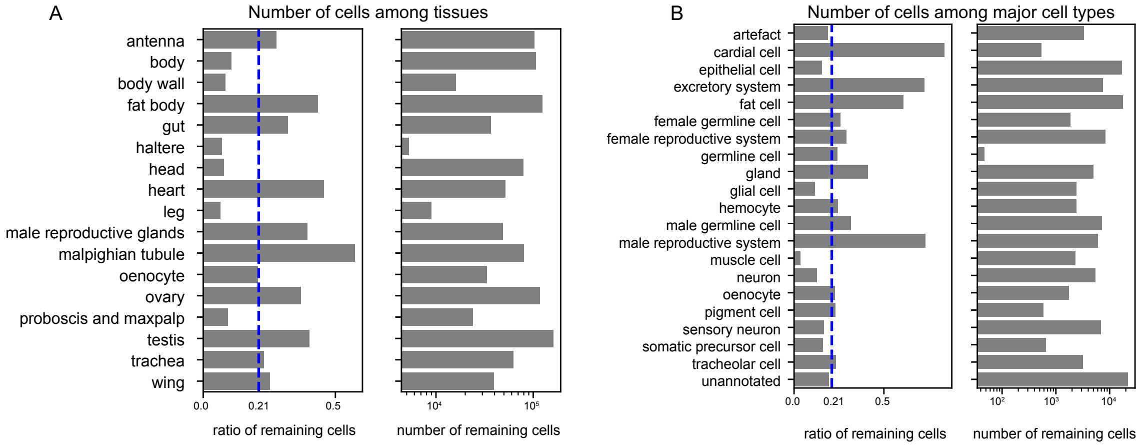


Figure S10. Ratio and number of remaining cells in different tissues (A) and major cell types (B) in the final filtered sub-expression matrix.

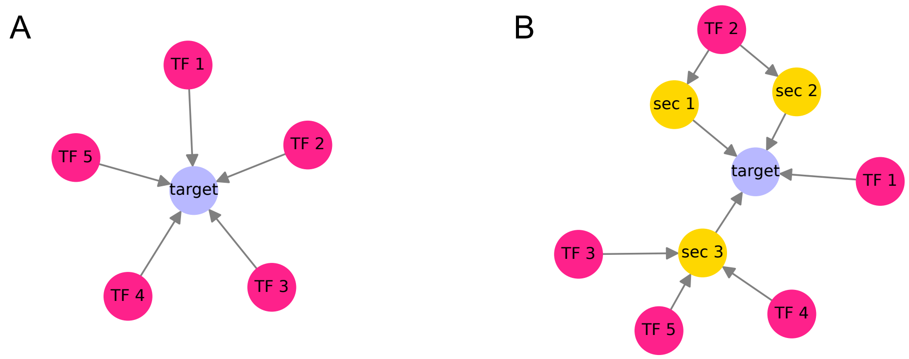


Figure S11. Simplified diagram of the transcriptional regulation network analysis illustrating number of direct transcription factors (TFs) and number of transcriptional regulation paths from transcription factors to target genes. In the diagram, TF i, {i=1,2,3,4,5}, represents five different major transcription factors, and sec i, {i=1,2,3}, represents three different secondary transcription factors. The target in (A) has five direct transcription factors. Number of transcriptional regulation paths of the five direct regulations is one. The target in (B) has four direct TFs, while the number of transcriptional paths from each of the five major and three secondary transcription factors, to the target are different. For example, the number of transcriptional paths from TF 1, TF 3, TF 4, TF 5, sec 1, sec 2, and sec 3, to the target is one, while the number of transcriptional paths from TF 2 to the target is two, which are TF 2 → sec 1 → target and TF 2 → sec 2 → target.

Table S1. Tissue specificity of top 31 regulons.

| FBgn | Symbol | Tissue-specificity (tau) | category | Enhanced_tissue |
|---|---|---|---|---|
| FBgn0033748 | vis | 0.9930756649506648 | top regulons | Testis |
| FBgn0001291 | Jra | 0.5013664911042334 | top regulons | |
| FBgn0033749 | achi | 0.8954960765344512 | top regulons | Testis |
| FBgn0261617 | nej | 0.6720055110651855 | top regulons | |
| FBgn0001147 | gsb-n | 0.9337617399901136 | top regulons | Testis |
| FBgn0003145 | prd | 0.99756546 | top regulons | AccessoryGland |
| FBgn0015239 | Hr78 | 0.7731994971901806 | top regulons | SalivaryGlandL |
| FBgn0022361 | Pur-alpha | 0.7051210235396731 | top regulons | GanglionM |
| FBgn0000577 | en | 0.9315734989648036 | top regulons | HindgutL |
| FBgn0039938 | Sox102F | 0.7224100544951354 | top regulons | |
| FBgn0001148 | gsb | 0.9191419141914192 | top regulons | |
| FBgn0000541 | E(bx) | 0.6738668386110713 | top regulons | Ovary |
| FBgn0000546 | EcR | 0.7285842878904781 | top regulons | |
| FBgn0040765 | luna | 0.45896657 | top regulons | |
| FBgn0004914 | Hnf4 | 0.6428282769967959 | top regulons | |
| FBgn0000015 | Abd-B | 0.8825035917296521 | top regulons | RectalPadM |
| FBgn0283521 | lola | 0.5759634727733687 | top regulons | |
| FBgn0287186 | scro | 0.9200625564084094 | top regulons | CropF |
| FBgn0014018 | Rel | 0.5327359020247745 | top regulons | |
| FBgn0004892 | sob | 0.8034557235421167 | top regulons | |
| FBgn0011648 | Mad | 0.53838805 | top regulons | |
| FBgn0003053 | peb | 0.8612348721837773 | top regulons | MidgutM |
| FBgn0038391 | GATAe | 0.91221741 | top regulons | MidgutM |
| FBgn0000028 | acj6 | 0.8873962429008301 | top regulons | Testis |
| FBgn0003507 | srp | 0.8657852029700768 | top regulons | |
| FBgn0086680 | vvl | 0.8125142466377934 | top regulons | |
| FBgn0005612 | Sox14 | 0.8957541938469867 | top regulons | MidgutL |
| FBgn0002521 | pho | 0.68972648 | top regulons | BrainF |
| FBgn0001235 | hth | 0.8555093839277169 | top regulons | VSp_ |
| FBgn0003345 | sd | 0.5243927542198435 | top regulons | |
| FBgn0032223 | GATAd | 0.5187347757558389 | top regulons | |

Table S2. The correlations between the expression levels of vis and other transcription factor genes involved in spermatogenesis. Different categories of these transcription factors are shown, including subunits in tMAC, tTAF and tPAF.

| FBgn | symbol | Slope | Intercept | R_squared | P_value | Padj | Category |
|---|---|---|---|---|---|---|---|
| FBgn0004372 | aly | 0.24154311 | 13.4966548 | 0.69380024 | 0.00011561 | 0.00736811 | tMAC |
| FBgn0034667 | comr | 0.10972264 | 4.78835423 | 0.62234965 | 0.00047256 | 0.01401405 | tMAC |
| FBgn0031715 | tomb | 0.21566761 | 24.8906125 | 0.58666646 | 0.00087045 | 0.01957998 | tMAC |
| FBgn0263979 | Caf1-55 | 0.22094126 | 20.7678064 | 0.49351196 | 0.00349521 | 0.04396605 | tMAC |
| FBgn0033770 | wuc | 0.45104349 | 57.4060075 | 0.58136829 | 0.00094905 | 0.0207891 | tMAC |
| FBgn0037751 | topi | 0.16505577 | 27.0866084 | 0.27651794 | 0.04407823 | 0.18842002 | tMAC |
| FBgn0034430 | mip40 | 0.22440535 | 100.45091 | 0.19636951 | 0.09805741 | 0.29915669 | tMAC |
| FBgn0040339 | MED22 | 0.16079919 | 47.1192731 | 0.4044936 | 0.01081438 | 0.0850909 | Mediator |
| FBgn0011569 | can | 0.19008282 | 11.9727518 | 0.57153833 | 0.00111122 | 0.02256831 | tTAF |
| FBgn0014342 | mia | 0.04705362 | 11.1175821 | 0.41533464 | 0.00950065 | 0.0782582 | tTAF |
| FBgn0002842 | sa | 0.50024672 | 67.9680452 | 0.48089026 | 0.00414363 | 0.0484734 | tTAF |
| FBgn0041103 | nht | 0.32554816 | 46.4865567 | 0.33082921 | 0.02488119 | 0.13670353 | tTAF |
| FBgn0031623 | Taf12L | 0.11789157 | 105.022433 | 0.03489106 | 0.50503758 | 0.73567946 | tTAF |
| FBgn0032366 | Taf9L | 0.36586992 | 39.0289905 | 0.76164096 | 2.18E-05 | 0.00442704 | tTAF |
| FBgn0027524 | Ski8 | 0.324944 | 48.6250631 | 0.53551891 | 0.00192753 | 0.03104453 | tPAF |
| FBgn0031388 | Paf1L | 0.53353025 | 54.9106705 | 0.63424663 | 0.00038083 | 0.01240157 | tPAF |
| FBgn0033865 | Cdc73L | 0.24136803 | 19.8813775 | 0.56309705 | 0.00126904 | 0.02446781 | tPAF |
| FBgn0034829 | Ctr9L | 0.31667005 | 7.24884356 | 0.70282227 | 9.47E-05 | 0.00670442 | tPAF |
| FBgn0038773 | Leo1L | 0.55623388 | 17.3634077 | 0.65190693 | 0.00027298 | 0.01046185 | tPAF |
| FBgn0039124 | tBRD-1 | 0.63492633 | 43.657439 | 0.66833903 | 0.00019733 | 0.00917014 | bromodomain |
| FBgn0034423 | tBRD-2 | 0.97996189 | 27.5141158 | 0.72643015 | 5.45E-05 | 0.00552863 | bromodomain |
| FBgn0000158 | bam | 0.02755438 | 2.78570761 | 0.46170718 | 0.00533001 | 0.05643735 | other |
| FBgn0016917 | Stat92E | 0.05093756 | 6.5792835 | 0.23651543 | 0.06603255 | 0.23783089 | other |
| FBgn0086758 | chinmo | 0.05437937 | 6.77619301 | 0.24475381 | 0.06081717 | 0.22673454 | other |
| FBgn0033749 | achi | 0.21113895 | 7.72575578 | 0.36903858 | 0.01630437 | 0.10758868 | paralog |
